# Muscle transcriptome analysis reveals molecular pathways and biomarkers involved in extreme ultimate pH and meat defect occurrence in chicken

**DOI:** 10.1101/101170

**Authors:** Stéphane Beauclercq, Christelle Hennequet-Antier, Christophe Praud, Estelle Godet, Anne Collin, Sophie Tesseraud, Sonia Métayer-Coustard, Marie Bourin, Marco Moroldo, Frédéric Martins, Sandrine Lagarrigue, Elisabeth Le Bihan-Duval, Cécile Berri

## Abstract

The processing ability and sensory quality of chicken breast meat are highly related to its ultimate pH (pHu), which is mainly determined by the amount of glycogen in the muscle at death. To unravel the molecular mechanisms underlying glycogen and meat pHu variations and to identify predictive biomarkers of these traits, a transcriptome profiling analysis was performed using an Agilent custom chicken 8×60K microarray. The breast muscle gene expression patterns were studied in two chicken lines experimentally selected for high (pHu+) and low (pHu-) pHu values of the breast meat. Across the 1,436 differentially expressed (DE) genes found between the two lines, many were involved in biological processes related to muscle development and remodelling and carbohydrate and energy metabolism. The functional analysis showed an intensive use of carbohydrate metabolism to produce energy in the pHu- line, while alternative catabolic pathways were solicited in the muscle of the pHu+ broilers, compromising their muscle development and integrity. After a validation step on a population of 278 broilers using microfluidic RT-qPCR, 20 genes were identified by partial least squares regression as good predictors of the pHu, opening new perspectives of screening broilers likely to present meat quality defects.

## Introduction

Poultry meat is known for its relative inexpensiveness and good nutritional quality. However, the competitiveness of the poultry industry is impacted by several quality defects such as the well-known PSE- and DFD-like syndromes, which are respectively related to low and high meat pH^1^,and the emerging white striping (WS)^2,3^ and wooden breast (WB)^4^ defects, whose incidence is currently growing at a rapid pace. PSE- and DFD syndromes are directly related to muscle glycogen storage at death, which is the main determinant of meat’s ultimate pH (pHu) in chicken^5^. Several recent studies also highlighted that lower glycogen storage in muscle predisposes breast meat to the WS and WB conditions^3,6^. The genetic and physiological control of muscle glycogen in avian species is still largely unknown compared to mammals in which several mutations have been identified in relation to muscle glycogen and meat quality variations. Indeed, several mutations of the γ3 regulatory subunit of AMPK have been shown to be responsible for glycogen accumulation in muscle^7,8^ and, in the case of the pig, low meat pHu and poor processing ability^9^. Studies performed in chicken have not yet identified major genes responsible of the pHu variation, suggesting a polygenic determinism of the trait^10,11^. Therefore, unravelling the biological processes and molecular pathways underlying glycogen variations in chicken muscle remains an important objective that would help to limit the occurrence of the main meat quality defects in poultry^7,8^. In this perspective, a divergent selection on the pHu of the breast *pectoralis major* muscle was carried out on a commercial line representative of the current broiler performances. After only five generations of divergent selection^12^, this experiment led to the creation of two lines, namely the pHu- and the pHu+ lines, which are respectively characterised by a very low and very high pHu. These two lines were recently fully characterised for their phenotypes related to meat quality attributes^12–14^ and for their serum and muscle metabolomics profiles^15^. The metabolomics approach allowed for the identification of 20 and 26 discriminating metabolites in the serum and muscle, respectively, and a set of 7 potential pertinent biomarkers of the pHu in the serum^15^. To gain a better understanding of the molecular pathways that underlie pHu variations in the breast meat of broilers, a chicken Custom 8×60K Gene Expression Agilent Microarray (Agilent Technologies, Santa Clara, CA, USA) was used to perform a transcriptomic analysis using the same individuals that had been previously selected for the metabolome profiling. The analysis of the transcriptome enabled the identification of a subset of potential muscle genetic biomarkers of pHu using a univariate filtered sparse partial least squares (fsPLS) approach^16^, whose predictive value was further tested on a large number of animals (n=278) by quantitative reverse transcription PCR (RT-qPCR) microfluidic arrays.

## Results

### Animals and breast *pectoralis major* muscle phenotypes

The transcriptome analysis was performed on 15 and 16 individuals from the pHu- and pHu+ lines, respectively. These individuals were chosen for their extreme low or high breast meat pHu values within each line. Their average phenotype values are described in Table 1. The individuals of the two lines showed a similar body weight, breast meat yield and abdominal fatness. However, they showed a difference in the breast meat pHu close to 0.8 pH units, corresponding to a 50 μM/g (lactate equivalent) difference in the muscle glycolytic potential, which is representative of the muscle glycogen content at death. The two populations were also characterised by great differences in terms of meat quality, especially the lightness, drip loss, and toughness, which were much higher in the pHu- line than in the pHu+ line, while the processing yield presented the opposite results (Table 1).

**Table 1.**
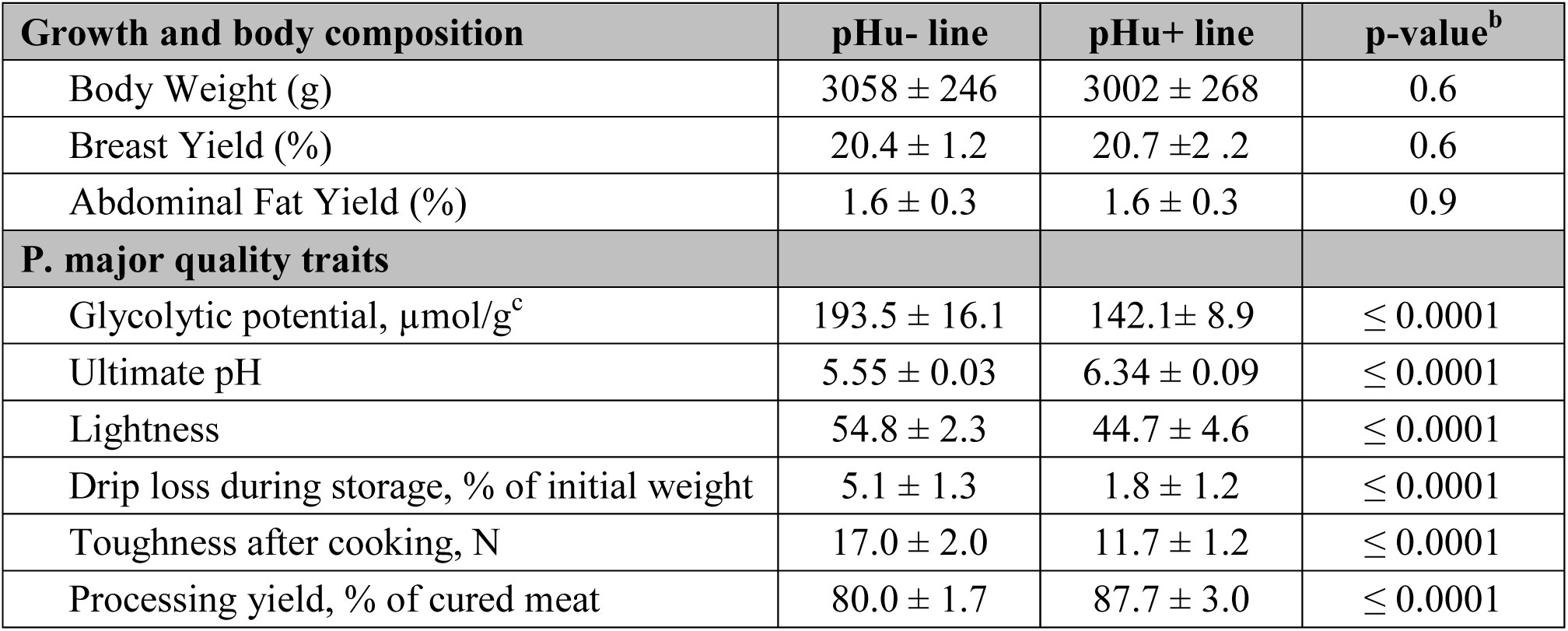
Body composition and *pectoralis major* muscle traits^a^ of the pHu- (n=15) and pHu+ (n=16) individuals. ^a^ Values are expressed as the mean ± SD; ^b^ Two-tailed Welch’s mean values equality t-test at 95% confidence level; ^c^ Measured at 15 min post-mortem. test at 95 % confidence level; ^c^ Measured at 15 min post-mortem.

| Growth and body composition | pHu- line | pHu+ line | p-value<sup>b</sup> |
| --- | --- | --- | --- |
| Body Weight (g) | 3058 ± 246 | 3002 ± 268 | 0.6 |
| Breast Yield (%) | 20.4 ± 1.2 | 20.7 ± 2.2 | 0.6 |
| Abdominal Fat Yield (%) | 1.6 ± 0.3 | 1.6 ± 0.3 | 0.9 |
| **P. major quality traits** |  |  |  |
| Glycolytic potential, $\mu\text{mol/g}^c$ | 193.5 ± 16.1 | 142.1 ± 8.9 | $\leq 0.0001$ |
| Ultimate pH | 5.55 ± 0.03 | 6.34 ± 0.09 | $\leq 0.0001$ |
| Lightness | 54.8 ± 2.3 | 44.7 ± 4.6 | $\leq 0.0001$ |
| Drip loss during storage, % of initial weight | 5.1 ± 1.3 | 1.8 ± 1.2 | $\leq 0.0001$ |
| Toughness after cooking, N | 17.0 ± 2.0 | 11.7 ± 1.2 | $\leq 0.0001$ |
| Processing yield, % of cured meat | 80.0 ± 1.7 | 87.7 ± 3.0 | $\leq 0.0001$ |

Muscle cross-sections from the two populations (pHu+ and pHu-) were further characterised using several histochemical stains. As expected, most of the pHu+ muscle fibres were depleted of glycogen despite their glycolytic metabolism (Fig. 1a and b). The muscle fibres of the pHu+ individuals were also characterised by a higher number of cells expressing the developmental embryonic and neonatal isoforms of the fast myosin heavy chain (Fig. 1e-h).

**Figure 1.**
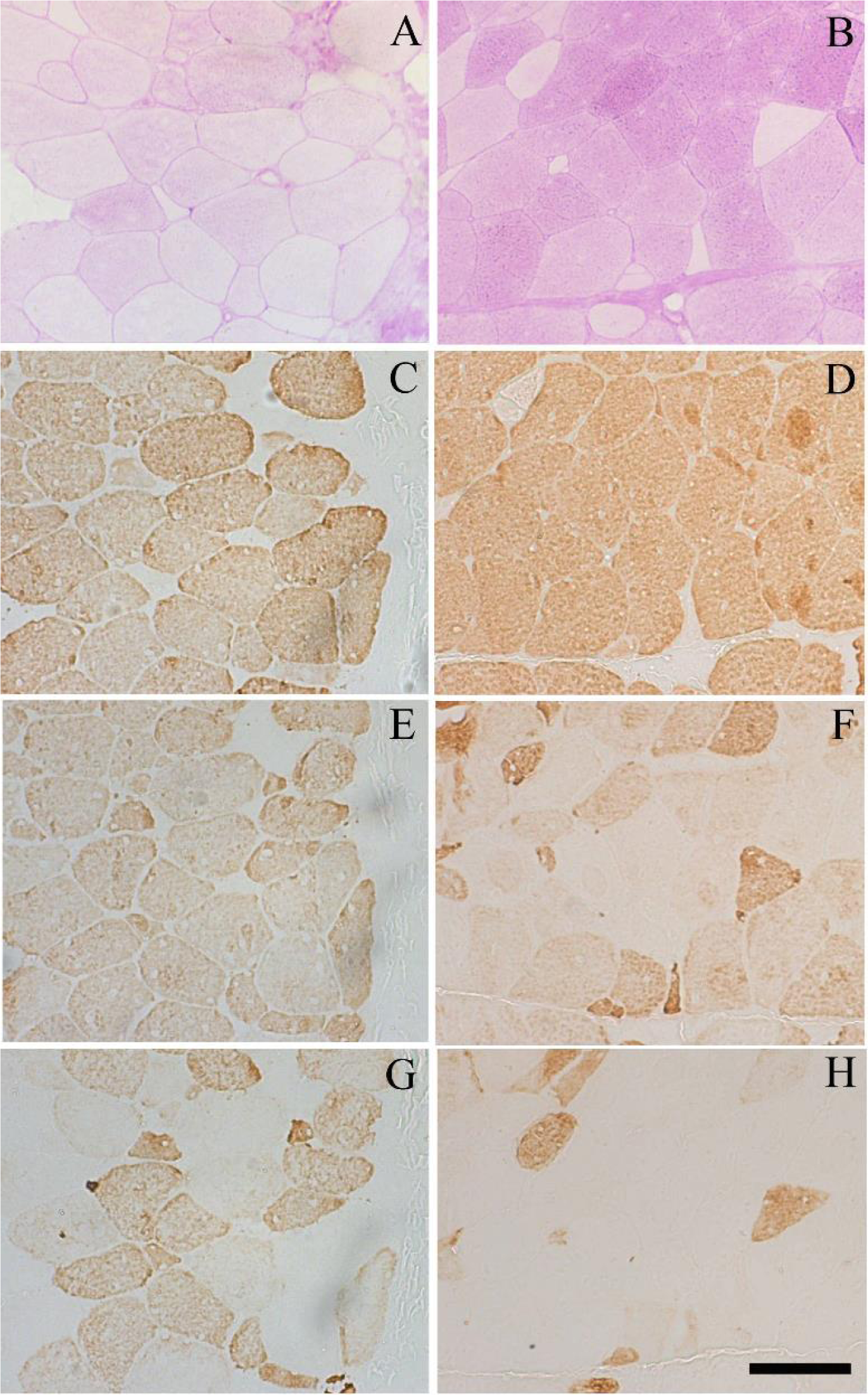
Serial light micrographs of the pHu+ (a, c, e, g) and pHu- (b, d, f, h) *pectoralis major* muscles. Periodic acid shift histochemical staining (**a, b**), immunohistochemical staining of the adult (**c, d**), both embryonic and neonatal (**e, f**), and neonatal (**g, h**) isoforms of the fast myosin heavy chain. Scale bar: 100 μm.

### Gene expression and functional annotation analyses

Among the 61,657 probes spotted on the array, 39,653 (64%) corresponded to genes that were expressed in the chicken *pectoralis major* muscle tissue. This corresponded to 15,789 unique genes, each of them being represented by a single probe after k-means clustering. Among these genes, 1,436 (approximately 10%) were considered statistically and biologically differentially expressed (DEGs) between the pHu- and pHu+ lines (adjusted p-value ≤ 0.05 and fold-change ≥ 1.2 or ≤ 0.8). A total of 850 genes were upregulated and 586 were downregulated when comparing the pHu- individuals to the pHu+ ones. As expected, the heat map obtained from the clustering of the DEGs (Fig. 2) showed a clear discrimination between the two groups characterised by the low or high pHu values. The range of the fold-change (FC) ran between 0.21–0.8 and 1.2–3.9 in the down- and upregulated gene groups, respectively. Lists of the top 10 down- and upregulated genes are provided as supplementary data (Supplementary Table S1 online).

**Figure 2.**
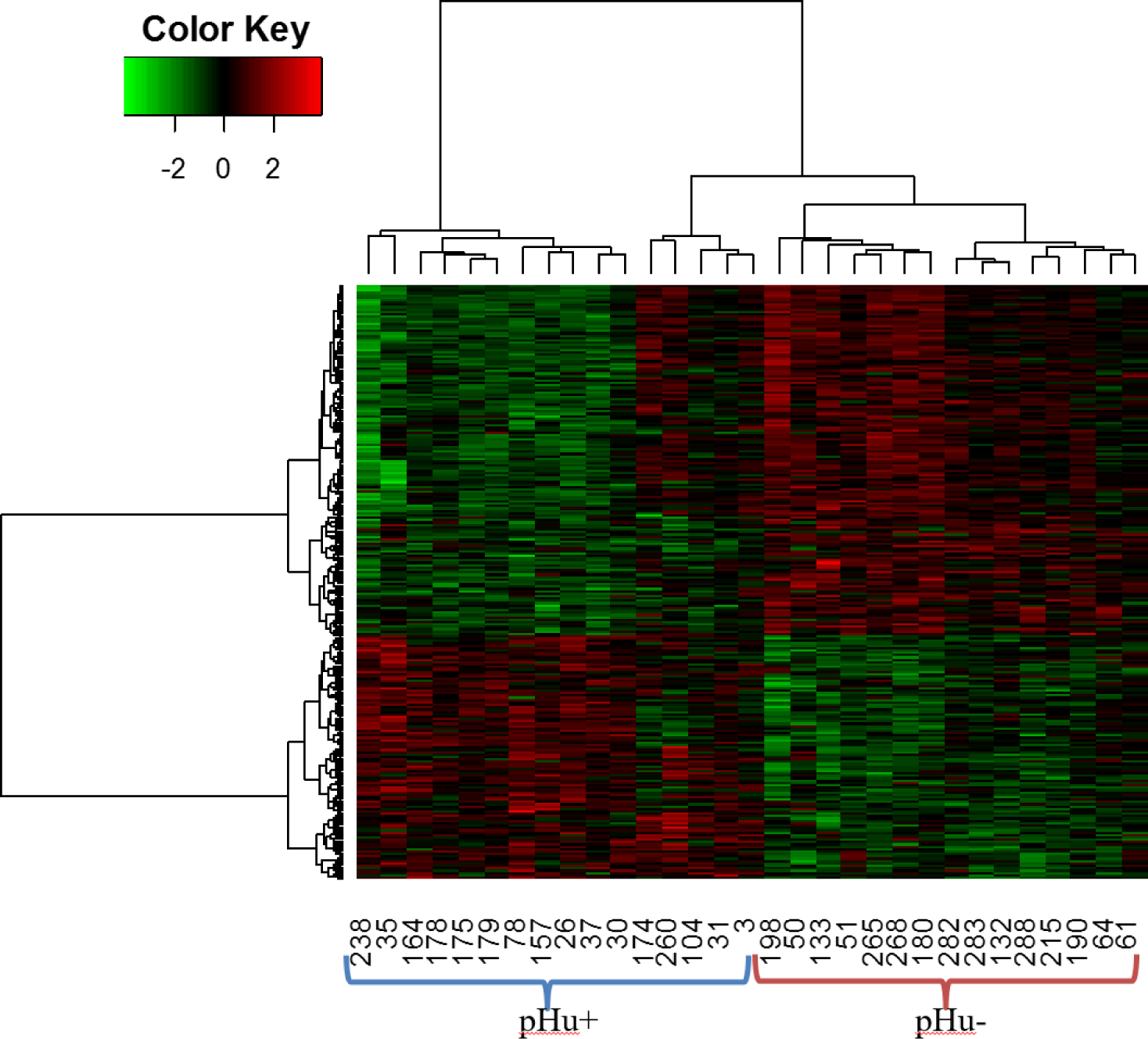
Heat map from the hierarchical clustering of differentially expressed genes between the pHu- and pHu+ individuals. The scaled expression by row (gene) is shown as a heat map and is reordered by a hierarchical clustering analysis (Pearson’s distance and Ward’s method) on both rows and columns.

Functional annotation was performed using an enrichment analysis with Fisher’s exact test. The over-representation of gene ontology (GO) terms within the group of DEGs was tested in the biological process ontology using the topGo R package, using the human orthology for collecting GO terms. Among the 1,436 DEGs, only 849 with human orthologues were mapped to Biological Pathway (BP) GO terms. The top 10 enriched BP GO terms were considered (Fig. 3). The most significantly enriched terms were canonical glycolysis with 53% of DEGs, muscle filament sliding (30%), wound healing (14%), muscle cell cellular homeostasis (40%), insulin-like growth factor receptor signalling pathway (31%), regulation of integrin-mediated signalling pathway (57%), gluconeogenesis (23%), cardiac muscle tissue morphogenesis (26%), nucleotide diphosphate metabolic process (29%), and modulation of synaptic transmission (17%). It is worth noting that within the processes linked to carbohydrates (canonical glycolysis and gluconeogenesis), most of the DEGs (90%) were overexpressed in the pHu- line.

**Figure 3.**
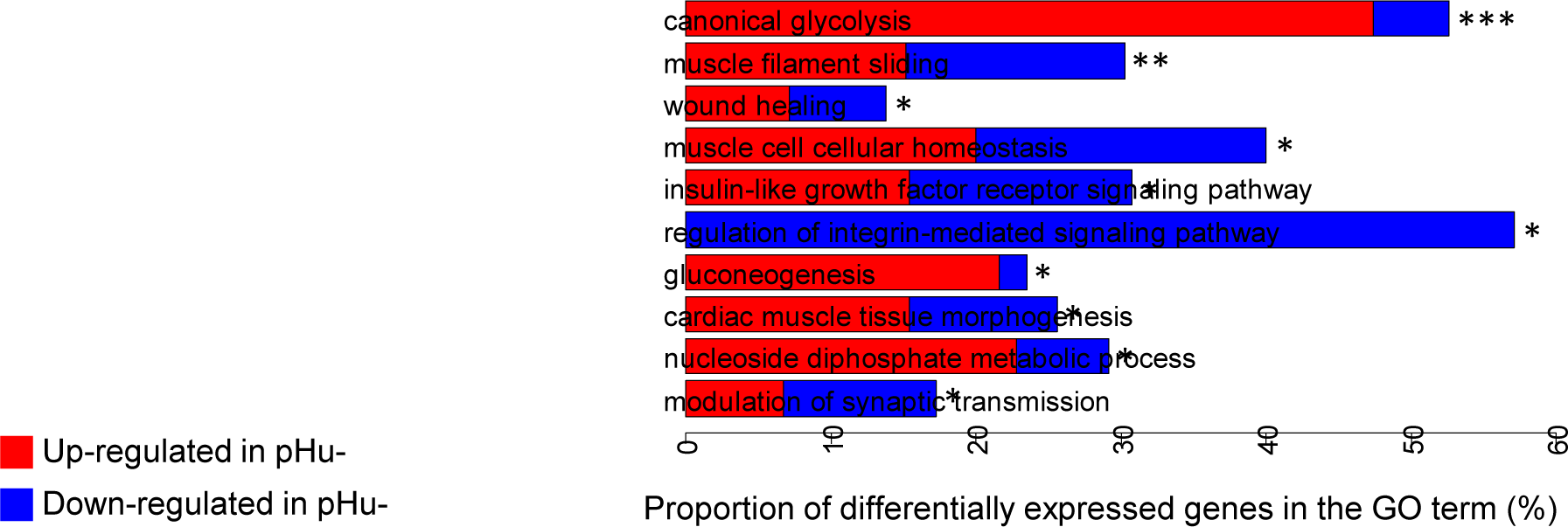
Top 10 enriched biological process GO terms. A functional enrichment analysis of the biological process GO terms was performed using the topGo R package on the genes that were differentially expressed in the breast *pectoralis major* muscle from the two lines of broilers divergently selected for their breast meat ultimate pH. The levels of significance are indicated by *** p-value ≤ 1×10^-5^, ** p-value ≤0.001, and * p-value ≤ 0.01.

### Selection of putative biomarkers of breast muscle pHu using the fsPLS method

A first step of filtering by dimension reduction was necessary before the sparse partial least squares (sPLS) analysis in order to limit overfitting and select the relevant genes predicting the quantitative pHu variable. This filtered sPLS model (fsPLS) underwent a cross-validation (CV) procedure. At each step of the CV, a differential analysis was performed and a sPLS model, including the DEGs that were filtered, was fitted with two components. From 672 to 1,000 DEGs were filtered by the differential analysis and were included in sPLS models with a sparsity criterion of 50 genes to explain the pHu variation and select the relevant genes. Only the first component of the sPLS models was informative, with an explanatory power R^2^ ranging from 0.77 to 0.87 and a predictive ability Q^2^ from 0.68 to 0.79. The robustness of the prediction accuracy was assessed using the mean square error of prediction (MSEP) estimated on the test samples. The MSEP ranged from 0.02 to 0.49, with an average of 0.23. A stability analysis, based on the frequency of genes selected in the fsPLS, was performed. A total of 134 genes were retained in at least one fsPLS model, while 15 genes were common at each of the 10 CV steps (Table 2). A total of 21 genes were kept in at least half of the fsPLS models and were retained for further validation on a large experimental population composed of 278 males and females derived from both the pHu+ and pHu- lines (Supplementary Table S2 online).

**Table 2.** List of the 15 genes kept in all 10 fsPLS models fitted to identify predictive biomarkers of chicken breast meat pHu.

| Gene symbol | Description | Associated biological process Go term |
| --- | --- | --- |
| ATP2B1 | ATPase, Ca<sup>++</sup> transporting, plasma membrane 1 | Calcium ion transmembrane transport |
| LSMEM1 (C7ORF53) | Leucine-rich single-pass membrane protein 1 |  |
| CDAN1 | Codanin 1 | Negative regulation of DNA replication, chromatin assembly, chromatin organization, ... |
| LOC100859584 (CR353014) | Gas2-likeprotein 1-like |  |
| LOC107052650 (CR387500) | Non coding RNA |  |
| ID3 | Inhibitor of DNA binding 3, HLH protein | Negative regulation of myoblast differentiation, response to wounding, positive regulation of apoptotic process, ... |
| MORN4 | MORN repeat containing 4 |  |
| MYLIP | Myosin regulatory light chain interacting protein | Positive regulation of protein catabolic process, protein ubiquitination involved in ubiquitin-dependent protein catabolic process, cholesterol homeostasis, ... |
| PCGF5 | Polycomb group ring finger 5 | Positive regulation of transcription from RNA polymerase ii promoter, transcription, DNA-templated |
| PPFIBP1 | PPFIA binding protein 1 | Cell adhesion |
| PRKCH | Protein kinase C eta | Protein phosphorylation, ... |
| SLC34A2 | Solute carrier family 34 member 2 | Sodium-dependent phosphate transport, cellular phosphate ion homeostasis, ... |
| THSD7B | Thrombospondin type 1 domain containing 7B |  |
| TNS1 | Tensin 1 | Fibroblast migration, cell-substrate junction assembly |

### Validation using microfluidic RT-qPCR of a subset of 48 DE genes revealed by the microarray analysis as potential biomarkers of the chicken breast meat pHu

Among the genes found to be differentially expressed between the pHu+ and pHu- individuals, 48 were selected for validation on a large population of 278 male and female broilers derived from both the pHu+ and pHu- lines. This population included the 31 extreme birds chosen for the transcriptome analysis, 57 contemporary individuals producing DFD meat (pHu > 6.1), 79 producing acid meat (pHu < 5.7), and 111 producing “normal” meat with a range of pHu values between 5.7 and 6.1. The genes under analysis included the 21 genes present in more than 50% of the pHu predictive fsPLS models described above, 6 genes showing a very high or low fold-change according to the microarray analysis, and 21 genes whose function is related to muscle carbohydrate and energy metabolism (Supp. Fig. S1, Table S4).

First, the fold-changes of DEGs obtained using microfluidic RT-qPCR were compared to the ratios obtained using microarrays on the subset of 31 individuals shared by both types of analyses (Fig. 4). Figure 4 reveals a high Pearson’s correlation coefficient between the log2 fold-change (pHu-/pHu+) of gene expression from the microarray and microfluidic technologies (0.84, p-value = 8.31×10^-14^). When considering the whole population, i.e., the 278 chickens derived from the pHu+ and pHu- lines, the differences in gene expression measured by RT-qPCR between the two animal groups was confirmed by an analysis of variance (p-value ≤ 0.05) for 18 of the 21 genes determined by the fsPLS models and only 9/28 of the other ones. The differentially expressed genes are highlighted in Supplementary Table S4.

**Figure 4.**
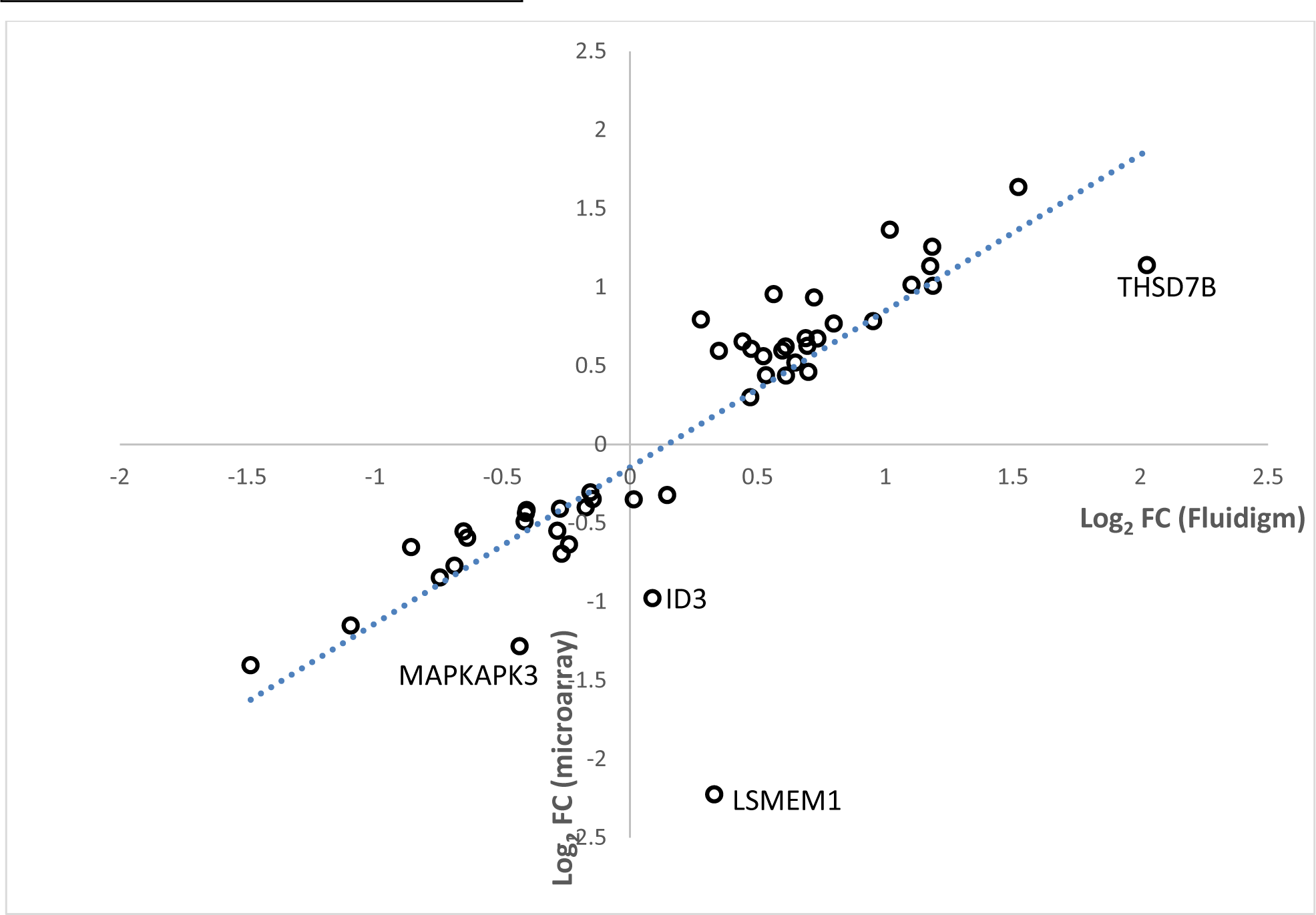
Correlation between the log2 fold-change (pHu-/pHu+) of gene expression from microarray and quantitative RT-PCR microfluidic technologies.

Second, a first PLS model with 48 variables (corresponding to the 48 genes whose expression was measured using microfluidic RT-qPCR) was fitted on the entire population of 278 individuals to explain the pHu variability. For predictive purposes, a parsimonious PLS model with the most relevant variables (variable important in projection score (VIP) ≥ 1) was subsequently fitted without lowering the predictive ability. This final PLS model included 20 genes and 3 components with a cumulative R^2^(cum) and Q^2^(cum) of 0.65 and 0.62, respectively (Fig. 5a). The model explained 65% of the observed variability of pHu with 35% for the first component. The scatterplot of the observed pHu vs. the predicted pHu revealed a strong linear relationship (Fig. 5b), and the root mean square error of estimation (RMSEE) was 0.16. This model included 14 genes from the fsPLS model (Supplementary Table S2 online), three showing extreme FCs according to the microarray analysis and three with functions related to carbohydrate or muscle metabolism (Figure 5). Loadings on the 3 components of the PLS model of the 20 genes kept in the model are provided in online Supplementary Table S3.

**Figure 5.**
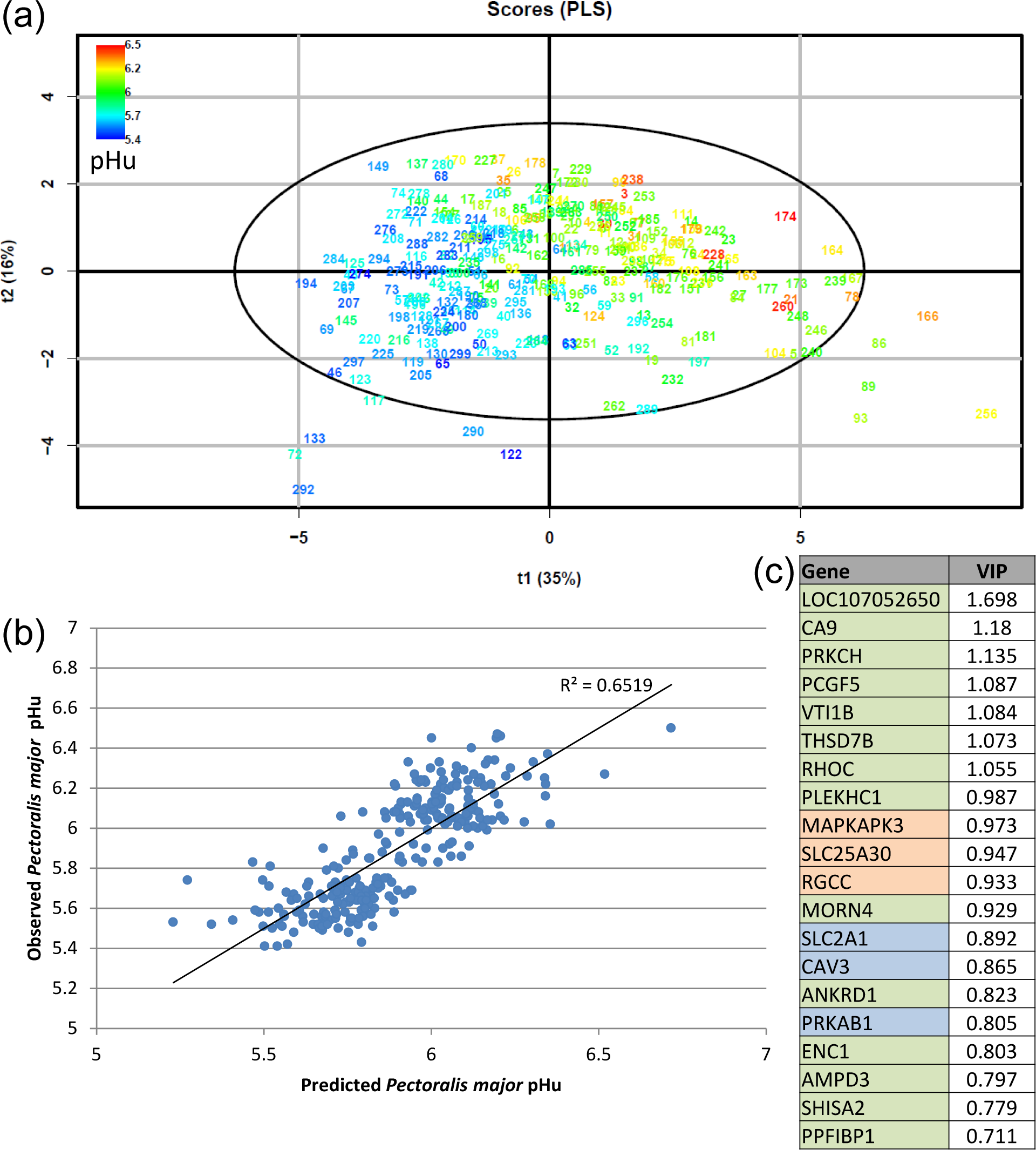
Final PLS model for the *pectoralis major* pHu prediction. The final PLS model was fitted on the mRNA expression of 20 genes (measured using RT-qPCR microfluidic technology) on a population of 278 male and female individuals from the pHu+ and pHu- lines in order to predict the *pectoralis major* pHu. (**a**) Score plot of the two first components (t1, t2) from the PLS model. Points correspond to chicken tags coloured according to their measured pHu value in the *pectoralis major* muscle. The cumulative explanatory and predictive performance characteristics of the model are R^2^(cum) = 0.65 and Q^2^(cum) = 0.62. The root mean square error of the estimation (RMSEE) was 0.16. (**b**) Scatterplot of the observed pHu vs. predicted pHu. (**c**) Variable Importance in Projection (VIP) of each variable (genes) in the final PLS model (the genes from the fsPLS model are highlighted in green, those with an extreme FC based on microarray analysis are in orange, and those whose function is directly related to carbohydrate and muscle metabolism are in blue).

## Discussion

In recent years, we have assisted in the incredible development of ‘foodomics’ coupled with high throughput techniques, aiming not only to improve the nutritional and gustative quality of food products but also to reduce waste due to product quality defects^17^. However, there is still a lack of understanding of the mechanisms underlying the variations in muscle pHu, which represents a major determinant of the sensory traits and processing ability of meat in poultry^18^.

To improve our knowledge of the genetic and physiological control of meat pHu, we recently developed two broiler lines (pHu+ and pHu-) that exhibit quite different muscle pHu values after 5 generations of divergent selection but show, at the same time, similar growth performance and body composition^12^. The between-line difference in the pHu values is related to significant differences in the muscle glycogen content and many other meat quality traits, including colour, water-holding capacity, texture, processing yield, as well as white striping occurrence^3,12–14^. Therefore, the pHu+ and pHu- lines constitute an excellent model to study the molecular pathways specifically involved in the control of poultry meat quality traits related to pHu and glycogen storage in muscle. The muscle transcriptome analysis described in the present study revealed that regulations related to carbohydrates and energetic pathways as well as to muscle remodelling were most impacted by the selection of meat based on pHu. The over-activation of glycolysis and gluconeogenesis pathways observed in the breast muscle of the pHu- line is directly related to its over-abundance of energy stored as muscle glycogen and ATP^15^. On the contrary, the lack of energy store that characterised the pHu+ muscles seems to be consistent with the overexpression of many genes involved in catabolic and muscle regeneration processes as well as in response to oxidative stress.

### Most genes involved in carbohydrate and energy metabolism are upregulated in the muscle of the pHu- line

The breast muscles of the pHu- line exhibited 36% more glycogen content compared to those of the pHu+ line. Therefore, it is expected that the metabolic pathways activated to produce energy differ between the two lines. The muscle transcriptome analysis showed that most of the genes involved in glycolysis were upregulated in the pHu- line, with gene expression increases that were between 24 and 70%. The only exception was aldolase C, which was downregulated (-29% in pHu-). Our previous observations showed that broilers from the pHu- line were characterised by slightly higher amounts of glucose in the serum and of glucose-6-phosphate (G-6-P) and fructose 1,6-bisphosphate in muscle when compared to those of the pHu+ line^15^. Therefore, this is consistent with the observation that the glucose-6-phosphate isomerase that transforms G-6-P to fructose-6-phosphate (F-6-P) during the first step of glycolysis and the phosphofructokinase that catalyses the conversion of F-6-P into fructose 1,6-bisphosphate (F1,6P) were overexpressed (+37 and +38%, respectively) in this line. Among the genes directly involved in carbohydrate metabolism, β-enolase-3 showed the highest fold-change (+70% in the pHu- line). A deficiency in this gene is associated with the glycogen storage disease type XIII and a decreased production of ATP^19^. Similarly, the glucose-6-phosphate transporter (SLC37A4), which is implicated in the glycogen storage disease type Ib^20^, was also overexpressed in the pHu- line (+45%). The role of this endoplasmic reticulum-bound transporter, coupled with glucose-6-phosphatase, is to maintain glucose homeostasis between meals. It is reported that a deficiency in one of these two proteins results in a phenotype of disturbed glucose homeostasis^21^.

It is worthy to note that the upregulation of most of the genes related to the glycolysis pathway is likely to increase the level of pyruvate entering the citric acid cycle, and thus, higher level of ATP are produced, as was observed with high-resolution^31^ P NMR^15^ in the muscle of pHu- animals. The overabundance of ATP observed in the pHu- muscle is in turn likely to enhance glycogen synthesis by glycogen synthase^22^.

The expression levels of several key regulators of glycogen turnover were also affected by selection of breast meat pHu (Fig. 6). Indeed, the protein phosphatase-1 regulatory subunit 3A (PPP1R3A), which binds glycogen with high affinity, activates glycogen synthase (GYS), and inhibits glycogen phosphorylase kinase (PHK) by dephosphorylation through the protein phosphatase-1 catalytic (PPP1C) subunit, was upregulated (+72%) in the pHu- line. Moreover, the regulatory β subunit of the phosphorylase kinase (PHKB) was also upregulated in pHu- muscle tissue (+60%). It has been reported that the overexpression of PPP1R3A and its catalytic subunit in glycogen-depleted cells leads to slightly higher GYS and lower glycogen phosphorylase (PYG) activity, resulting in increased glycogen content^23^. Moreover, there are variants of PPP1R3A that impair glycogen synthesis and reduce muscle glycogen content in humans and mice^24^. The low expression of PPP1R3A in the pHu+ line is therefore consistent with the low level of muscle glycogen that characterised this line. The selection of the breast meat pHu also affected the expression of several subunits of the AMP-activated protein kinase (AMPK) complex, another key regulator of glycogen turnover. The AMPK complex consists of one α catalytic subunit and two non-catalytic subunits, β and γ, and each is represented by several isoforms. The β subunit contains a binding domain to glycogen and a protein-protein interaction domain for the formation of the heterotrimeric AMPK complex by the binding of the α and γ subunits. The γ subunit has four repeat motifs CBS (cystathionine-β-synthase) involved in the binding of AMP and ATP. Quite interestingly, we observed an overexpression of the β2 (+35%) at the expense of the β1 (-34%) subunit in the pHu- muscle when compared to the pHu+ muscle. A similar regulation of the β1 subunit of the AMPK complex was observed in another chicken model. In the lean and fat lines that were originally selected for low and high abdominal fatness but that also diverge for breast muscle glycogen content (fat > lean) and meat quality, β1 was downregulated in the fat line that exhibited the highest glycogen content in muscle^11^. The γ3 subunit of AMPK was also overexpressed (+54%) in the muscle of the pHu- compared to the pHu+ broilers. As mentioned above, the γ subunits of the AMPK complex act as energy sensors in the muscle cell as they contain binding domains to AMP and ATP^25^. Several mutations in the γ2 subunit are associated with glycogen accumulation in the human heart (Wolff-Parkinson-White syndrome)^20^, while a mutation of the γ3 subunit results in increased glycogen in pig muscle and production of acid meat with a poor processing ability^9^. Some PRKAG3 SNPs were identified in chicken leading to variations in their meat quality, but their potential biological roles remain unknown^26^. However, there is no evidence that an upregulation of this gene at the transcript level is a sufficient condition to induce glycogen accumulation in muscle^8^.

**Figure 6.**
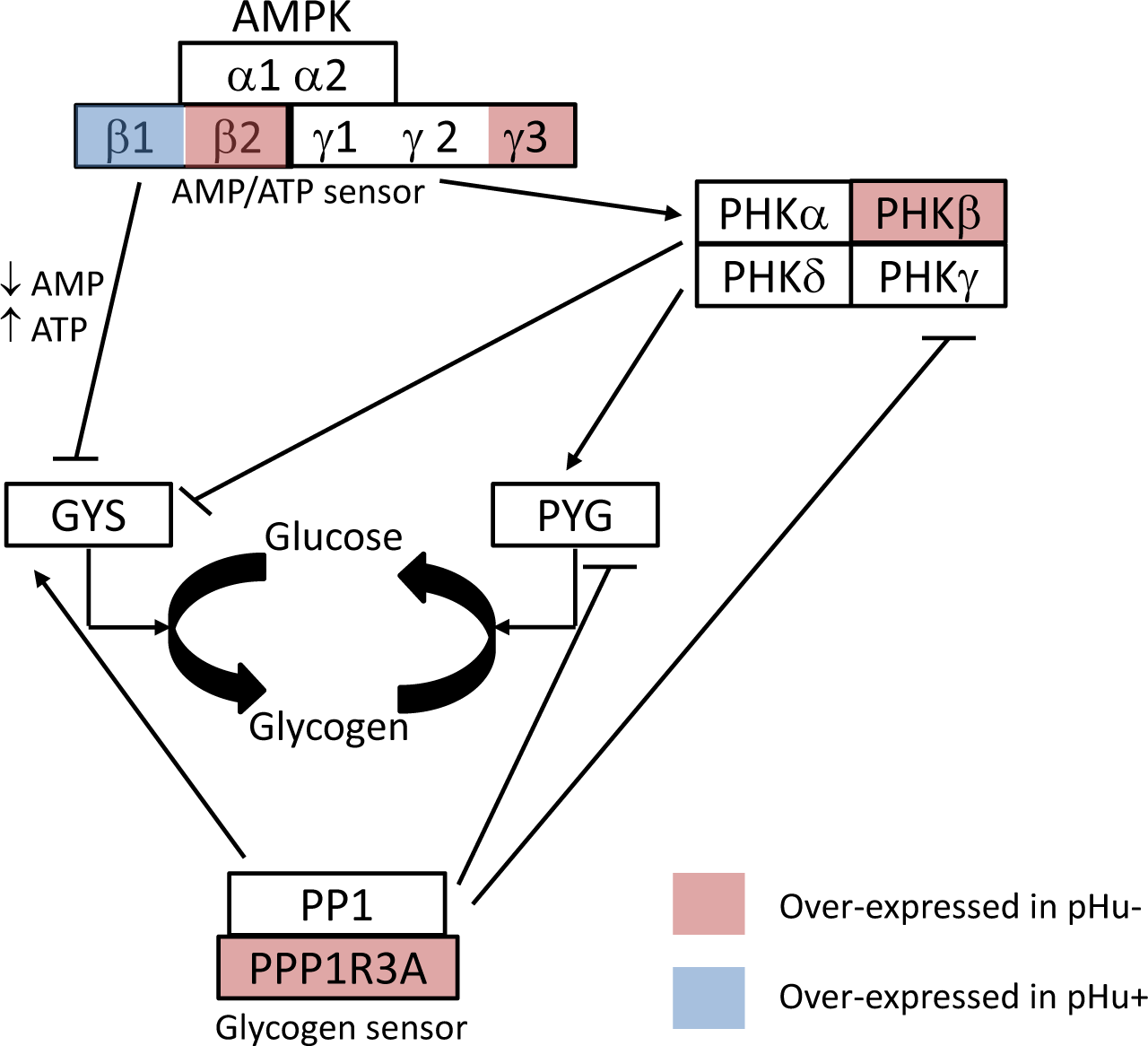
Molecular actors of glycogen turnover. Function of the studied enzymes in the regulation of muscle glycogen metabolism. AMPK = adenosine monophosphate-activated protein kinase; PHK = phosphorylase kinase; GYS = glycogen synthase; PYG = glycogen phosphorylase; PP1 = protein phosphatase-1; PPP1R3A = protein phosphatase-1 regulatory subunit 3A.

Beyond the genes directly involved in the regulation of glycogen turnover, several other genes that were overexpressed in the pHu- compared to the pHu+ line may influence glycogen storage in muscle. This is the case for the gene encoding phosphodiesterase 3B (PDE3B, +23%) whose activation by insulin may induce antiglycogenolytic effects^27^ or the one encoding mitochondrial creatine kinase (CKMT2, +59%), which is responsible for the transfer of high-energy phosphate from mitochondria to creatine. This observation can therefore be clearly linked to the higher content of phosphocreatine (+36%) and in general to the higher energetic status that characterised the pHu- compared to the pHu+ muscles^15^.

Our recent metabolomics study highlighted that muscles of the pHu+ line solicit more intense oxidative pathways, such as lipid β-oxidation and ketogenic amino acid degradation, to produce energy and compensate for the lack of energy due to carbohydrates and glycolysis^15^. At the transcript level, this results in the regulation of two genes in the pHu+ muscle. 3- hydroxymethyl-3-methylglutaryl-CoA lyase (HMGCL), which catalyses the final step of leucine catabolism and ketone body formation in the mitochondria, and acetyl-CoA acetyltransferase-2 (ACAT2), which transforms the acetoacetyl-CoA (resulting from β-oxidation or degradation of ketogenic amino acids) into acetyl-CoA^28^, were upregulated (+21% and +36%, respectively) in pHu+ muscle. Finally, there are several genes encoding nudix hydrolases that are overexpressed in the pHu- muscle compared to the pHu+ muscle, including NUDT7 (+49%)^29^, NUDT12 (+27%)^30^, and NUDT19 (+38%)^31^. Nudix hydrolases hydrolyse a wide range of organic pyrophosphates, including nucleoside di- and triphosphates, dinucleoside and diphosphoinositol polyphosphates, nucleotide sugars and RNA caps^32^. They are implicated in the elimination of oxidised CoA and the regulation of CoA and acyl-CoA levels in the peroxisome in response to metabolic demand^29^ but also in the regulation of β-oxidation^33^, which may differ between lines^15^.

### Genes involved in protein breakdown, muscle remodelling, and response to oxidative stress were upregulated in the pHu+ line

Interestingly, a gene encoding m-calpain (CAPN2) was upregulated (+62%) in the pHu+ muscle. Calpains are Ca^2+^-activated cysteine proteases involved in the calcium-dependant proteolytic system. Their overexpression was reported in cases of muscular dystrophy^34^ or in the muscle of chicken subjected to a transitory dietary lysine Deficiency^35^. They are known to catalyse a limited proteolysis of proteins involved in cytoskeletal modelling (e.g., desmin and vimentin) or signal transduction and also to play a role in regeneration processes^34,36^. In the pHu+ muscle, the upregulation of the two genes encoding desmin and vimentin, which belong to the muscle intermediate filaments^37^ (+45% and +30%, respectively), also supports the hypothesis of more intense muscle regeneration in this line. Moreover, the overexpression of the interferon related developmental regulator 1 (IFRD1, +44% in pHu+) and of the Leiomodin-2 (LMOD2, +103%), which mediates nucleation of actin filaments and thereby promotes actin polymerisation, could also be a good indicator of myoblast differentiation and skeletal muscle regeneration^38^. Caveolin 3 (CAV3), which plays a key role in muscle development and physiological processes as maintenance of plasma membrane integrity, vesicular trafficking and signal transduction, was also upregulated (+46% in pHu+)^39^.

Histological observations made in this study highlighted a greater expression of embryonic and neonatal myosin heavy chain isoforms in the pHu+ muscle than in the pHu- muscle of 6-week-old chickens, which is likely the sign of a stronger muscle fibre regeneration process^40,41^. However, it is difficult to relate the difference observed at the protein level in muscle cross-sections to those observed at the gene level in the microarray. Indeed, several DNA probes match genes encoding fast myosin heavy chains (MHC), but the high degree of similarities existing between the RNA sequences makes the association between the probes and antibodies used for histological analysis difficult to determine. Nevertheless, among the 12 fast myosin isotypes present on the microarray, three were overexpressed (+32% for MHC2, +48% for MHC6, +35% for MHC9) while one was under-expressed (MHC1, −48%) in the pHu- line when compared to the pHu+ line. This strongly supports our hypothesis that the muscle fibre regeneration process is accompanied by a switching within the fast myosin isotypes. All of these observations are consistent with recent data that suggest that muscle glycogen depletion is related to meat defects in which degeneration and regeneration processes occur. Indeed, the occurrence and severity of white striping are higher in the pHu+ than in the pHu- line^3^. Moreover, wooden breast-affected muscles exhibit a much lower glycogen content that is associated with increased oxidative stress, elevated protein levels, muscle degradation, altered glucose utilisation and redox homeostasis when compared with samples from unaffected birds^6^.

Hypoxanthine and xanthine, which are both products of purine degradation and may be markers of oxidative stress^42^, were the most differential serum metabolites between the pHu+ and pHu- lines (pHu+ > pHu-)^15^. Logically, the gene encoding xanthine dehydrogenase (XDH), which successively catalyses the oxidation of hypoxanthine to xanthine and the one of xanthine to urate^43^, tended to be overexpressed in the pHu+ muscle (+55%, FDR adjusted p-value < 0.1). Genes implicated in the cellular oxidative homeostasis, such as glutathione peroxidases (GPX3, +21% and GPX8, +37%) or glutathione reductase (GSR, +22%), were also upregulated in the pHu+ muscles. These observations are consistent with the observation that higher levels of antioxidant molecules present in the pHu+ line^15^ likely prevent the negative cellular effects of oxidative stress in this line.

### Identification of biomarkers predictive of breast meat pHu

In addition to the integrated view of the transcriptional changes occurring in muscle in response to selection, our analysis was also an opportunity to identify potential biomarkers for this trait and more generally for chicken meat quality. Contrary to what is generally practiced in transcriptomic studies, we applied k-means clustering to average the signal of probes targeting the same gene and showing a similar regulation. This prevents the loss of information by combining differential and non-differential probes targeting the same gene. Indeed, mutations, alternative splicing or early transcription ending may lead to differential regulation within probes belonging to the same gene, and this diversity of response is of high interest in a biomarker discovery approach. An sPLS model is a relevant approach to select the most predictive genes for pHu variation from a gene expression microarray. Although sPLS usually performs better than PLS, it is not able to overcome the overfitting issue by itself^16^. Therefore, and as suggested by Le Floch *et al*^16^, dimension reduction and 10-fold cross-validation steps were performed to limit overfitting and extract a robust association between gene expression and pHu. This resulted in 10 fsPLS models based on one informative component including 50 genes. Among them, 15 genes were systematically kept in all 10 models, which identifies them as being the most robust genes to predict breast meat pHu in chicken. It is relevant to observe that these genes included a majority of genes implicated in muscle organisation but not directly involved in carbohydrate metabolism. Interestingly, the expression of Ankyrin Repeat Domain 1 (ANKRD1), which is implicated in lipid metabolic process and sarcomere organisation, was overexpressed (+71%) in the pHu+ line and is also shown to be positively correlated to pork meat pHu^44^. Among this set of 15 genes, there were two genes whose expression was the most differential among all of the identified DEGs, and these included LOC107052650 (adjusted p-value FDR = 4.11×10^-9^) and LSMEM1 (adjusted p-value FDR = 1.31×10^-7^). It is striking that they are both localised in a narrow region (26,908K - 26,659K) of chromosome 1 (Gallus gallus 5.0 assembly) and that quite close, in the 26,361K - 26,388K region, is a major regulator of glycogen turnover, PPP1R3A, whose expression was significantly higher in pHu- muscle compared to pHu+ muscle. Altogether, these observations indicate that this region of the genome is important for the genetic determinism of breast meat pHu in chicken.

To go further in the identification of biomarkers of breast meat pHu, we decided to enlarge the list of potential pHu biomarkers and to evaluate their predictive ability on a large population of 278 chickens covering a pHu range from 5.41 to 6.50. Therefore, we considered 21 genes issued from the fsPLS (kept in more than 50% of the predictive models) and 27 additional genes, including 6 exhibiting very high or low fold-changes between the pHu- and pHu+ lines, as well as 21 whose function is directly related to muscle carbohydrate and energy metabolism. A parsimonious PLS model was fitted on the expression of 20 genes among the 48 analysed by microfluidic RT-qPCR on the population of 278 chickens. The explicative and predictive abilities were good (R^2^ = 0.65, Q^2^ = 0.62), and the root mean square error of estimation was surprisingly low (16%). Concretely, this model allowed us to correctly classify 74% of the muscles exhibiting a normal value of pHu, i.e., those between 5.7 and 6.1, within the population of 278 broilers evaluated. Furthermore, it is worthy to note that 70% of the biomarkers kept in the model were issued from the fsPLS analysis, which highlights the robustness of combining both filtering (as differential analysis) and sparsity in the PLS approaches.

Any further development of predictive molecular tools would, however, require the validation of the predictive potential of the model on independent populations. The identification of this set of genes should contribute to advancing the understanding of the genetic control of this trait but also to better target nutritional or husbandry factors that regulate the storage of glycogen in the muscle and more generally the quality of breast meat in chicken.

In conclusion, the transcriptomics approach developed here provided an integrative view of the gene regulation that underlies the muscle capacity to store glycogen in chicken and highlighted several genes as predictive markers of chicken breast meat ultimate pH and quality defects. It clearly showed that muscles prone to produce acid meat are characterised by an overactivation of carbohydrate metabolism and highlighted an adaptive response of glycogen-depleted muscles that induces several catabolic, oxidative and cellular repair processes also involved in the setting up of white striping and wooden breast conditions. Therefore, biological markers identified through the comparison of the pHu+ and pHu- lines could be of great interest for poultry producers due to the fact that a lack of carbohydrate storage in chicken muscle appears to be a predisposing factor or an indicator of sensitivity to such myopathies whose increasing incidence may compromise the competitiveness and consumer acceptability of the broiler industry.

## Methods

### Animals and sample collection

The present study was carried out on chickens belonging to the sixth generation of two lines divergently selected for high (pHu+) or low (pHu-) *pectoralis major* muscle pHu values^12^. The Ethics Committee of Val de Loire for Animal Experimentation approved all of the animal care and procedures used in this experiment (program 00880.02). The birds were reared and slaughtered at 6 weeks of age at the PEAT experimental unit (INRA, Centre Val de Loire, Nouzilly, France). The broilers received ad libitum food and water until 8 h before slaughtering. Fifteen minutes after slaughter, samples of *pectoralis major* were collected, immediately snap frozen in liquid nitrogen and stored at −80°C until RNA extraction. The pHu of the *pectoralis major* muscle was measured a day after slaughtering using a pH meter by direct insertion of the electrode into the thickest part of the muscle. A sub-sample of 16 pHu- and 16 pHu+ male broilers was selected for further analysis from the entire population, exhibiting extremely low (≤5.6) and high (≥6.2) breast muscle pHu values, respectively.

### Histochemical traits

Ten-micrometre thick *pectoralis major* muscle cross-sections were used to perform periodic acid-Schiff (PAS) staining and immunohistochemistry. For the PAS staining, muscle cross-sections were fixed in Carnoy’s fixative and were incubated for 5 minutes in periodic acid 1% and 90 minutes in Schiff’s reagent (Sigma-Aldrich, Saint-Quentin Fallavier, France) before washing in tap water. For immunohistochemistry, muscle cross-sections were first incubated in 10% goat serum for 30 minutes and were then incubated for 1 hour with primary MF14, B103 and 2E9 antibodies (1/30). The MF14 antibody, developed by Donald A. Fischman, and the B103 and 2E9 antibodies, developed by E. Bandman, were obtained from the Developmental Studies Hybridoma Bank, created by the NICHD of the NIH and maintained at The University of Iowa, Department of Biology, Iowa City, IA 52242. The antibodies were detected using the Vectastain ABC kit (Vector Laboratories, Burlingame, CA, USA) followed by a 5-minute DAB (Sigma-Aldrich) incubation. The slides were mounted in Canada balsam (Merck KGaA, Darmstadt, Germany) after dehydration.

### RNA isolation

Total RNA extraction was performed using the RNeasy Mini Kit (Qiagen, Valencia, CA, USA) on *pectoralis major* muscle samples that were ground in liquid nitrogen. Residual genomic DNA was removed by DNase I treatment (Qiagen, Valencia, CA, USA). RNA concentrations were measured using a NanoDrop ND-1000 spectrophotometer (Thermo Fisher Scientific, Waltham, MA, USA), and their integrity was assessed using RNA 6000 Nano chips (Agilent Technologies, Santa Clara, CA, USA) run on a Bioanalyzer 2100 (Agilent Technologies).

### Microarray data production and statistical analyses

#### Labelling and hybridisation

Muscle transcriptional profiling was performed using an 8×60K Agilent custom platform developed by the PEGASE Genetics and Genomics team (INRA, UMR1348, Rennes, France). The platform is referenced under the GEO reference GPL20588. Thirty-two samples were processed, corresponding to 16 pHu+ and 16 pHu- individuals. One pHu- sample was considered an outlier after the principal component analysis and was thus removed from the analysis. All of the steps of the RNA labelling and microarray processing were performed by the CRB GADIE facility (INRA, UMR GABI, Jouy-en-Josas, France, http://crb-gadie.inra.fr/) as described in Jacquier *et al.*^45^. The microarray data were submitted to the Gene Expression Omnibus (GEO) microarray database (accession number GSE89268).

#### Differential analysis

The single channel microarray data were analysed using the R/Bioconductor software package Limma (Linear Models for Microarray Data)^46^. The signals of all of the probes were log2 transformed and were subsequently normalised by median centring of each array. The differential analysis was performed using a t-test on each probe, and the p-value was adjusted for multiple testing by the Benjamini-Hochberg method to control the False Discovery Rate (FDR)^47,48^. The difference in the expression between the two lines (pHu- *vs.* pHu+) was measured using the log2 transformation of the fold-change (lfc). Probes with an adjusted p-value < 0.05 and a |lfc| ≥ 0.26 were considered as statistically and biologically differentially expressed between the pHu- and pHu+ lines. In the case of several oligonucleotides mapping the same gene, a representative probe was defined using k-means clustering with the algorithm of Hartigan and Wong^49^. Basically, the representative probe was defined by the mean lfc of the probes, and the maximal adjusted p-values of the best cluster of the partition on probe data sets was defined by the coordinates (lfc, −log_10_ (adjusted p-value)) in the plane.

The scaled expression of the differentially expressed genes (DEGs) by row was displayed in a cluster heat map to identify clusters of genes and clusters of samples with a similar profile. The colour gradient was set to green for the lowest expression value in the heat map and to red for the highest expression value. Both the rows and columns were reordered to correspond to the hierarchical clustering results based on the Pearson correlation distance and Ward's minimum variance method. As a convention, the fold-changes of genes (ratio pHu- /pHu+) were only indicated if they were differentially expressed between the pHu+ and pHu- with an adjusted p-value ≤ 0.05.

#### Functional annotation

The biological interpretation of expressional data was performed using the topGo R/Bioconductor package (version 2.22.0) and the Human annotation database R/Bioconductor (org.Hs.eg.db version 3.2.3) ^50^. The over-representation of GO terms in the biological process ontology (BP) was tested with Fisher’s exact test and the elim algorithm within the group of DEGs compared to all of the expressed genes on the array.

#### Filtering and sparse Partial Least Squares (fsPLS) model

Sparse Partial Least Squares (sPLS) regression was applied to select the relevant genes predicting the quantitative variable pHu, including a step of regularisation based on L1 penalisation using the mixOmics R package (version 5.2.0) ^51^. To deal with the high number of variables and overfitting, a two-step approach fsPLS combining univariate filtering (i.e., differential analysis) followed by a sPLS model was performed, as described by Le Floch *et al.* ^16^. The robustness of the prediction accuracy was obtained using a cross-validation (CV) scheme. First, a 10-fold CV scheme was constructed to provide 10 training sets for gene expression and pHu of approximately the same size (26 or 27 samples) and with an almost balance between pHu- and pHu+. At each fold, a filter of at most 1000 of the best ranked DEGs (with |lfc| ≥ 0.26 and an adjusted p-value < 0.05) was obtained by differential analysis on the training gene expression set and consequently a new training set filtered for gene expression. Then, a sPLS model with two components, which were linear combinations of 50 arbitrary relevant genes, was fitted on each filtered training set and was evaluated on a corresponding test set (samples left out). To assess the explanatory power and the predictive accuracy of each fsPLS model, we estimated the R^2^ as a measure of the goodness of fit and Q^2^ as a measure of the predictive ability of the model using a 10-fold cross-validation on the training set and the MSEP on test sets. A stability analysis, based on the frequency of the genes retained in fsPLS models, was performed. A schematic representation of the fsPLS statistical method applied for pHu biomarker discovery is provided as online supplementary figure S1.

### Quantitative RT-PCR assay and the PLS model

Forty-eight genes, differentially expressed in the *pectoralis major* muscle between the pHu+ and pHu- lines, were selected for further analysis by RT-qPCR on 278 male and female chickens issued from the same populations (which include the 31 individuals used for microarray analysis). Total RNAs were extracted from *pectoralis major* muscle samples using RNA NOW (Ozyme, St Quentin en Yvelines, France). From each *pectoralis major* muscle sample, 10 μg of total RNA was reverse-transcribed using RNase H^-^ MMLV reverse transcriptase (Superscript II, Invitrogen, Illkirch, France) and random primers (Promega, Charbonnières les Bains, France).

The set of genes studied by RT-qPCR included (i) 21 genes present in more than 50% of the pHu predictive fsPLS models; (ii) 21 genes whose function was related to muscle energy metabolism; and (iii) 6 genes exhibiting high or low fold-changes according to the microarray differential analysis. Primers targeting those genes were designed with Primer3 version 4.0.0 (Supplementary Table S4 online)^52,53^. The corresponding amplicons were analysed by gel electrophoresis and were subsequently sequenced. Afterwards, the levels of expression of these genes in the *pectoralis major* muscle were quantified using a Fluidigm Biomark microfluidic platform (Fluidigm, South San Francisco, CA, USA). Two types of references were included for this type of analysis: the housekeeping gene CDK6 to normalise the Ct values of each target gene, and a mix of chicken *pectoralis major* muscle cDNAs as a calibrator sample. As previously described^54^, the calculation of the expression levels was based on the PCR efficiency and the threshold cycle (CT) deviation of target cDNAs versus the calibrator cDNAs, according to the equation proposed by Pfaffl^55^.

First, the RT-qPCR data were used to confirm the expression results obtained from the microarray by calculating the Pearson’s correlation between the log2 fold-change obtained by microarray and RT-qPCR using the *cor.test* R function.

Second, the pHu values from the 278 chickens population were fitted from the expression of the 48 genes measured by the RT-qPCR assay with a PLS model using the ropls Bioconductor R package^56^. The first three components of the model were considered to evaluate the prediction by their explicative (R^2^), predictive ability (Q^2^), and root mean square error of estimation (RMSEE) computed by a 7-fold cross-validation. To identify the most predictive genes, the variables with a variable importance in projection score (VIP) ≥ 1 were kept in a more parsimonious final model.

## Acknowledgements

We thank the staff of the Poultry Breeding Facilities (INRA, UE1295 Pôle d’Expérimentation Avicole de Tours, F-37380 Nouzilly, France) for producing the birds and the Avian Research Unit (INRA, UR83 Recherches Avicoles, F-37380 Nouzilly, France) for their valuable technical assistance. We also thank the StatOmique group for the valuable discussion on multivariate statistical methods. This study was supported by the French Ministry of Agriculture through the RFI CASDAR program no. 1309 OPTIVIANDE. S.B. was supported by a grant from the INRA and the French Ministry of Agriculture.

## Author information

### Contributions

S.B. drafted the first version of the manuscript under the supervision of CB. C.B. supervised the study. S.B. performed the gene expression, statistical, and functional analyses. C.H.A. supervised the statistical analyses and carried out the fsPLS model. C.P. carried out the histological characterisation of muscles. S.B., E.G, S.M.C., A.C., and S.T. designed and tested the oligonucleotide primers used for quantitative RT-PCR analysis. E.G. and S.B. performed RNA extraction. M.M. performed the microarray hybridisation. F.M. performed the high-throughput RT-qPCR. SL designed the ID *GPL20588* chicken Custom 8×60K Gene Expression Agilent Microarray platform. C.B., E.L.B.D., and M.B. contributed to the experimental design and organised the phenotype data collection. C.B., C.H.A, E.L.B.D., S.T., A.C. and S.M.C helped to draft the manuscript. All of the authors read and approved the final manuscript.

## Additional Information

### Supplementary information

**Table S1**, Top 10 up- and downregulated genes between the pHu- and pHu+ muscles. **Table S2**, Loadings of the fsPLS models. **Table S3**, Loadings of the three components of the parsimonious validation PLS model based on gene expression measured by RT-qPCR. **Figure S1**, Schematic representation of the statistical procedure implemented for pHu biomarker discovery. **Table S4**, oligonucleotide sequences of the primers used for mRNA quantification by RT-qPCR analysis.

### Accession GEO codes

GSE89268

